# Modeling toes contributes to realistic stance knee mechanics in three-dimensional predictive simulations of walking

**DOI:** 10.1101/2021.08.13.456292

**Authors:** Antoine Falisse, Maarten Afschrift, Friedl De Groote

## Abstract

Physics-based predictive simulations have been shown to capture many salient features of human walking. Yet they often fail to produce realistic stance knee mechanics and terminal stance ankle plantarflexion. While the influence of the performance criterion on the predicted walking pattern has been previously studied, the influence of the mechanics has been less explored. Here, we investigated the influence of two mechanical assumptions on the predicted walking pattern: the complexity of the foot segment and the stiffness of the Achilles tendon. We found, through three-dimensional muscle-driven predictive simulations of walking, that modeling the toes and metatarsophalangeal joints, and thus using two-segment instead of single-segment foot models, contributed to robustly eliciting physiological stance knee flexion angles, knee extension torques, and knee extensor activity. Yet modeling toe joints did not improve ankle kinematics, nor did decreasing the Achilles tendon stiffness. The lack of predicted terminal stance ankle plantarflexion thereby remains an open question. Overall, this simulation study shows that not only the performance criterion but also mechanical assumptions affect predictive simulations of walking. Improving the realism of predictive simulations is required for their application in clinical contexts. Here, we suggest that using complex models is needed to yield such realism.

## Introduction

Predictive simulations have advanced our understanding of gait neuromechanics but also revealed gaps in our current knowledge [1]. Motivated by the observation that humans select temporospatial walking features (e.g., step frequency [2] and stride length [3]) that minimize metabolic energy per distance traveled, predictive simulations of walking commonly rely on the assumption of performance optimization. *De novo* walking patterns can therefore be generated by solving for muscle controls that optimize a performance criterion based on a model of musculoskeletal dynamics with task-level constraints such as walking speed and periodicity. This approach has produced walking patterns that capture many salient features of human gait, indicating that performance optimization and musculoskeletal dynamics are important determinants of gait mechanics. Yet several gait features have remained hard to capture. Predicting knee flexion during stance along with physiologically plausible knee extension torques and activation patterns of the knee extensors (vasti) has been challenging. Another gait feature that has been hard to simulate, but has received far less attention, is ankle plantarflexion during terminal stance. While the role of the performance criterion has been explored, it remains unclear how musculoskeletal modeling assumptions contribute to the lack of stance knee flexion and terminal stance ankle plantarflexion in simulations of walking.

The performance criterion underlying predictive simulations of walking has been shown to influence the degree of knee flexion during stance in two-dimensional (2D) walking simulations. Yet it has been difficult to identify a criterion that robustly generates such flexed knee patterns. Ackermann and van den Bogert [4] found that stance knee flexion was influenced by the cost function in muscle-driven 2D simulations of walking. They compared energy-like cost functions, sum of muscle-volume-scaled activations to the power 1-4, and fatigue-like cost functions, sum of activations to the power 2-10, and found that energy-like cost functions resulted in straight stance knee patterns with low knee extensor activations, whereas fatigue-like cost functions resulted in more realistic knee flexion angles and knee extensor activations during stance. However, these observations might not be robust against other modeling choices as they were confirmed by some (e.g., [5]) but not all future predictive simulation studies of walking. For example, our recent 2D muscle-driven simulations of walking based on a similarly complex model lacked stance knee flexion although we solved for muscle controls that minimized a fatigue-like cost function (i.e., muscle activations cubed) [6]. Further, Koelewijn et al. [7] obtained realistic stance knee flexion, nevertheless in the absence of knee extension torques and knee extensor activations, when optimizing for metabolic energy.

Finding a performance criterion that produces realistic stance knee flexion has also been challenging when using more complex three-dimensional (3D) musculoskeletal models. Anderson and Pandy [8] simulated a 3D walking motion by solving for controls that minimized metabolic energy while enforcing knee flexion during stance by imposing joint kinematics at the initial time instant of the simulation, chosen to be mid-stance, to match experimental data. Miller [9] found that metabolic energy models influenced the simulated walking pattern, with most models predicting some, albeit less than experimentally observed, knee flexion during stance. All of the simulated walking patterns underestimated ankle plantarflexion in late stance. In recent work, we found that minimizing a cost function that penalized both metabolic energy and muscle activity yielded more realistic walking patterns than minimizing a cost function that penalized either metabolic energy or muscle activity [10]. Yet our simulations underestimated stance knee flexion, knee extension torques, and knee extensor activations as well as terminal stance ankle plantarflexion. Hence, control objectives other than reducing metabolic cost or fatigue as well as musculoskeletal modeling assumptions might also play a role in shaping stance knee and ankle mechanics.

Here, we demonstrate that musculoskeletal mechanics influences knee flexion during stance. We first focused on the foot, which is commonly modeled as a single-segment rigid body despite the clear extension of the metatarsophalangeal joints during walking. We found, through 3D simulations of walking, that a two-segment foot model that allowed movement between the toes and the rest of the foot led to more accurate predictions of knee joint kinematics and kinetics than a single-segment foot model. Yet modeling toe joints did not improve ankle kinematics. We therefore also explored the influence of the Achilles tendon stiffness on the simulated walking pattern. Achilles tendon compliance has been suggested to improve gait efficiency both by allowing for storage and release of energy throughout the gait cycle and through its influence on the operating length and velocity of the plantar flexors [11–15]. Importantly, it is the interaction between ankle kinematics and Achilles tendon properties that determines plantarflexor operating length and velocity, and thereby efficiency. Although both ankle and knee kinematics were sensitive to Achilles tendon stiffness, varying Achilles tendon stiffness within a physiological range did not improve the realism of the predicted walking patterns.

## Methods

The modeling and simulation workflow is described in detail in previous work [6,10]. Code and data will be made available at https://simtk.org/projects/3dpredictsim_toes and https://github.com/antoinefalisse/3dpredictsim (the code and data are available at https://github.com/antoinefalisse/predictsim_mtp while the paper is in review).

### Musculoskeletal model

We used an OpenSim musculoskeletal model with 31 degrees of freedom (DoFs) [16,17] (pelvis-to-ground: 6 DoFs, hip: 3 DoFs, knee: 1 DoF, ankle: 1 DoF, subtalar: 1 DoF, metatarsophalangeal-toe: 1 DoF, lumbar: 3 DoFs, shoulder: 3 DoFs, and elbow: 1 DoF), 92 muscles actuating the lower limb and lumbar joints, eight ideal torque motors actuating the shoulder and elbow joints, and six contact spheres per foot. To increase computational speed, we fixed the moving knee flexion axis of the generic model to its anatomical reference position [18]. We added passive stiffness (exponential) and damping (linear) to the lower limb and lumbar joints, henceforth referred to as passive torques, to model ligaments and other passive structures [8].

We adjusted the orientation of the toe joint coordinate frame of the generic OpenSim model to be aligned with that of its parent frame, namely the calcaneus. We found that an orientation of the toe joint axis perpendicular to the sagittal plane in the anatomical position better reproduced the movement of the toes during walking as compared to the generic toe joint orientation. The toe joints had no active actuation. On top of the aforementioned passive torques, we added a linear rotational spring with a stiffness of 25 Nm rad^−1^ [19] and a damper with a damping coefficient of 0.4 Nm s rad^−1^ for the toe joints. We used higher damping for the torque-driven toe joints as compared to the muscle-driven joints for which there is also damping at the muscle level. We used higher stiffness since there was no active actuation.

We used Raasch’s model [20,21] to describe muscle excitation-activation coupling and a Hill-type muscle model [22,23] to describe muscle-tendon interaction and the dependence of muscle force on fiber length and velocity. We modeled skeletal motion with Newtonian rigid body dynamics and smooth approximations of compliant Hunt-Crossley foot-ground contacts [10,17,24,25]. We described the dynamics of the ideal torque motors using a linear first-order approximation of a time delay. To increase computational speed, we defined muscle-tendon lengths, velocities, and moment arms as a polynomial function of joint positions and velocities. More details about the musculoskeletal model can be found in previous work [10].

### Experimental data

We used experimental data for comparison with simulation outcomes as well as to provide some of the bounds and initial guesses of the predictive simulations [10]. We collected data (marker coordinates, ground reaction forces, and electromyography) from one healthy female adult (mass: 62 kg, height: 1.70 m). The subject was instructed to walk over the ground at a self-selected speed. The average walking speed was 1.33 ± 0.06 m s^−1^. We processed the experimental data with OpenSim 4.2 [17]. The musculoskeletal model was scaled to the subject’s anthropometry based on marker information from a standing calibration trial. Joint kinematics were calculated based on marker coordinates through inverse kinematics. Joint kinetics were calculated based on joint kinematics and ground reaction forces. Electromyography was processed by band-pass filtering (second order dual-pass Butterworth filter, 20-400Hz), rectification, and low-pass filtering (second order dual-pass Butterworth filter, 10Hz).

### Predictive simulation framework

We formulated predictive simulations of walking as optimal control problems [6,10]. We identified muscle excitations and gait cycle duration that minimized a cost function subject to constraints describing muscle and skeleton dynamics, imposing left-right symmetry, preventing limb collision, and prescribing gait speed (distance traveled by the pelvis divided by gait cycle duration).

Our cost function included metabolic energy rate, muscle activity, joint accelerations, passive torques, and excitations of the ideal torque motors at the arm joints, all terms squared:

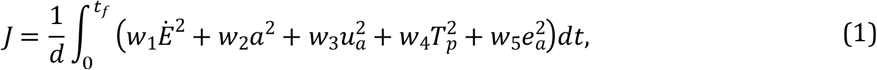

where *d* is distance traveled by the pelvis in the forward direction, *t*_*f*_ is half gait cycle duration, *a* are muscle activations, *u*_*a*_ are accelerations of the lower limb and lumbar joint coordinates, *T*_*p*_ are passive torques, *e*_*a*_ are excitations of the ideal torque motors driving the shoulder and elbow joints, *t* is time, and *w*_1―5_ are weight factors. We modeled metabolic energy rate 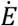 using a smooth approximation of the phenomenological model described by Bhargava et al. [26]. To avoid singular arcs [27], we appended a penalty function with the remaining controls to the cost function. More details about the optimal control problem formulation can be found in previous work [10].

We formulated our problems in Python 3.7 using CasADi 3.5 [28], applied direct collocation using a third order Radau quadrature collocation scheme with 100 mesh intervals per half gait cycle (the dynamic equations were therefore satisfied at 300 collocation points), used algorithmic differentiation to compute derivatives [6], and solved the resulting nonlinear programming problem with IPOPT [29] using a desired convergence tolerance of 10^−4^ (all other settings were kept to default). We generated our simulations from two initial guesses and selected results with the lowest optimal cost values (i.e., integral of equation 1 with optimal values). The first initial guess was a ‘hot-start’ based on experimental walking data, whereas the second initial guess was a ‘cold-start’ that consisted of a static, bilaterally symmetrical standing posture that translated forward over the simulation duration. Muscle states and controls were set to constant values for both initial guesses.

### Influence of the mass distribution and vertical location of the contact spheres

We investigated the effect of two main changes between the model used in the present study, henceforth referred to as new model, and the model used in our previous study [10], henceforth referred to as old model. The first change concerns the model mass distribution. In our previous study, we used a generic (i.e., pre-scaling) model with a heavier torso (34.2 kg) as compared to that of the generic model of the present study (26.8 kg). All other parameters, except inertia that scales with mass, were identical between both generic models. Since we scaled both models to the same subject mass, we obtained two scaled models with different mass distribution; the new model having a lighter torso but heavier legs as compared to the old model. The second change concerns the vertical location of the contact spheres. In our previous study, we used the foot-ground contact configuration from Lin et al. [30]. However, our predictions had an offset in the vertical position of the pelvis when compared to experimental data. In the present study, we therefore manually adjusted the vertical location of the contact spheres, moving them higher up (about 1 cm) to reduce that offset. We evaluated the sensitivity of the predicted walking pattern to those two changes.

### Influence of the toe joints

We evaluated the influence of incorporating toe joints in the model on the predicted walking pattern. To this aim, we compared predictive simulations of walking produced with the musculoskeletal model described above to simulations produced with the same model but with locked toe joints. In practice, for that model, we replaced the hinge toe joints by weld joints, thereby locking the toes in their neutral position. We performed this comparison for the old model (i.e., heavier torso and lighter legs) with low contact spheres as well as for the new model (i.e., lighter torso and heavier legs) with high contact spheres.

### Influence of the Achilles tendon stiffness

We evaluated the influence of the Achilles tendon stiffness on the predicted walking pattern. To this aim, we compared predictive simulations produced with Achilles tendon stiffness ranging from 30 to 100%, by increments of 10%, of the generic value (*k* = 35 [23]). In more detail, we adjusted the variable *k* for the triceps surae muscles (gastrocnemius lateralis and medialis, and soleus) in the following equation describing the tendon force-length relationship:

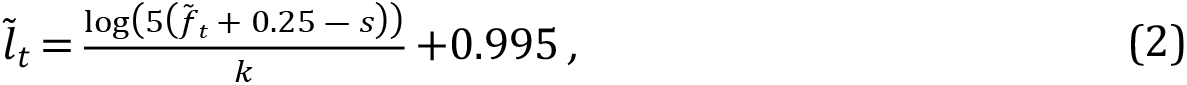

where 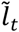 is the normalized tendon length, 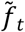 is the normalized tendon force, and *s* is a scaling parameter that shifts the tendon force-length curve as a function of the tendon stiffness *k* to enforce that the tendon force produced when the normalized tendon length equals the tendon slack length is independent of the tendon stiffness. We used the new model with toes for those simulations.

### Influence of the convergence tolerance and mesh density

We tested the sensitivity of the results to the number of mesh intervals by comparing walking patterns predicted with 50, 75, 100 (default), and 125 mesh intervals per gait cycle. Similarly, we evaluated the effect of the convergence tolerance by comparing the results produced with a convergence tolerance of 10^−4^ (default), 10^−5^, and 10^−6^. For both analyses, all numerical settings other than the one being studied were set to their IPOPT default.

## Results

### Influence of the mass distribution and vertical location of the contact spheres

Adjusting the model mass distribution had a small effect on knee kinematics and kinetics, barely any effect on ankle kinematics and kinetics, and a small effect on muscle activations (Fig 1). More specifically, the new model with a lighter torso and heavier legs produced slightly larger knee flexion angles, knee extension torques, and knee extensor (vasti) activations during stance. Moving the contact spheres higher up further increased knee flexion angles, knee extension torques, and knee extensor activations during stance. The mass distribution and contact location had little influence on the predicted metabolic cost of transport (between 3.89 and 3.93 J kg^−1^ m^−1^; Table 1), which is slightly above reported experimental measurements (3.35 ± 0.25 J kg^−1^ m^−1^ [9]).

**Table 1:** Influence of modeling choices on the metabolic cost of transport (COT).

|  | Without toe joints |  |  | With toe joints |  |
| --- | --- | --- | --- | --- | --- |
|  | Old model +<br>low contact<br>spheres | New model +<br>low contact<br>spheres | New model +<br>high contact<br>spheres | Old model +<br>low contact<br>spheres | New model +<br>high contact<br>spheres |
| COT<br>(J kg<sup>-1</sup> m<sup>-1</sup>) | 3.89 | 3.90 | 3.93 | 4.18 | 4.18 |

The new model has a lighter torso but heavier legs as compared to the old model. The high contact spheres are about 1 cm higher than the low contact spheres.

**Fig 1.**
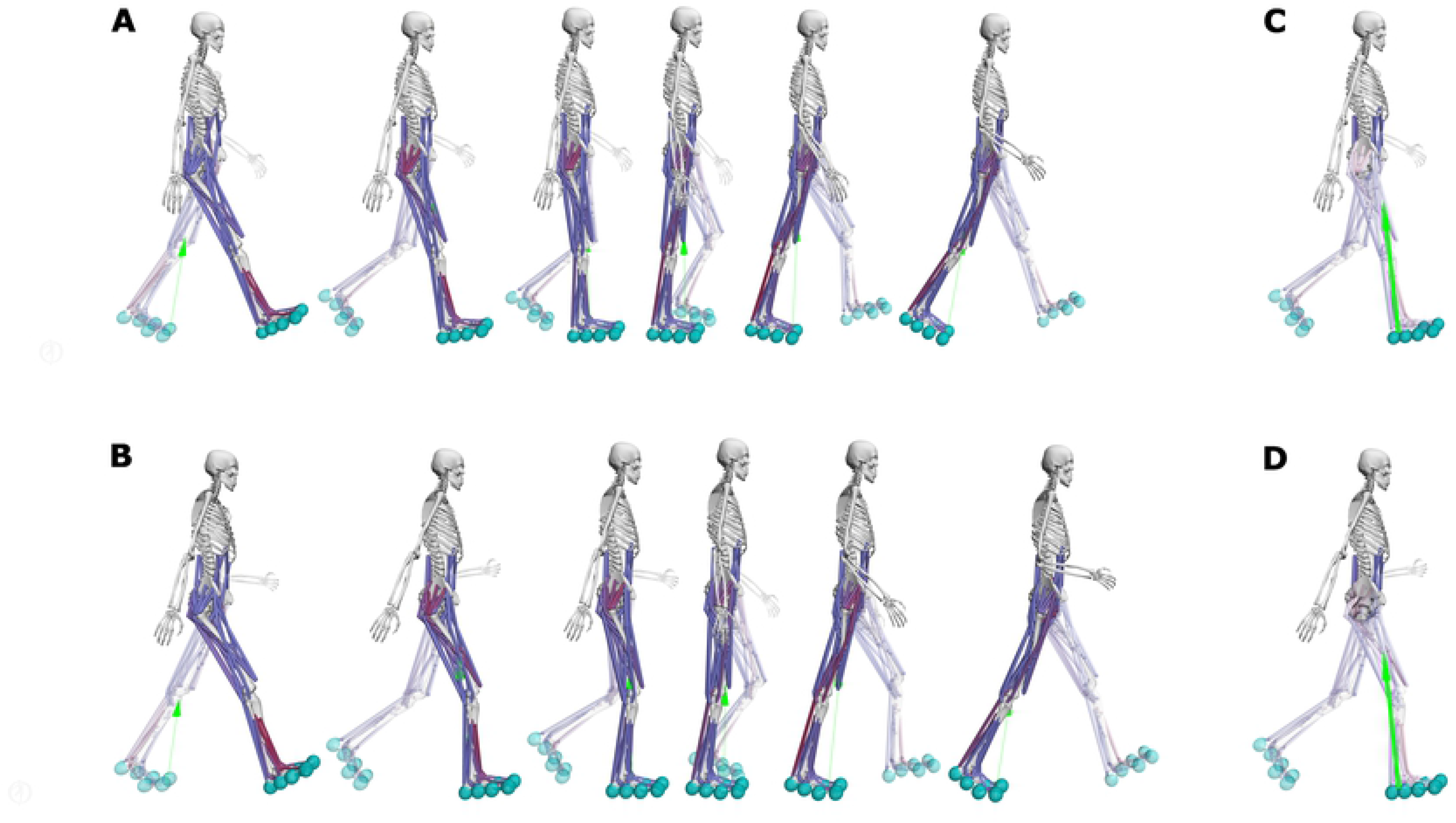
Influence of modeling choices on predicted gait pattern. Predicted knee and ankle kinematics (**A**) and kinetics (**B**), muscle activations (**C**), and ground reaction forces (**D**) with the different models. Experimental data (shaded areas) are shown as mean ± 2 standard deviations. The experimental electromyography data were normalized to peak activations of the new model with high contact spheres and with toes (dashdot orange curve). The old and new models have the same mass but different mass distribution: the new model having a lighter torso but heavier legs as compared to the old model. The difference in the vertical position of the contact spheres (about 1 cm) is illustrated in (**E**), and an example toe extension is depicted in (**F**). Results for all joints and muscles are shown in S1-3 Figs.

**Fig 2.**
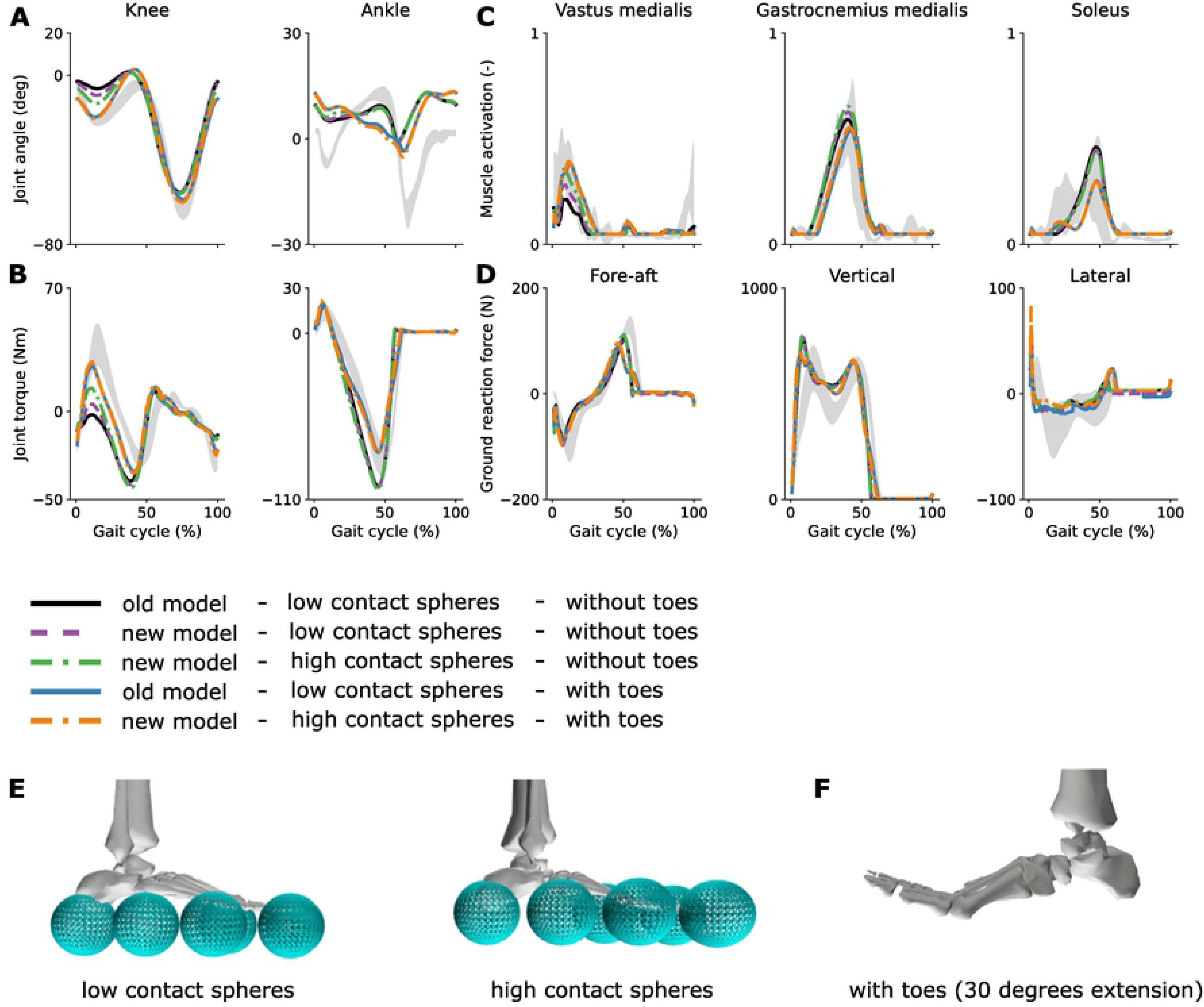
Snapshots of the predicted half gait cycles. Predicted half gait cycle with the old model with low contact spheres and without toes (**A**; solid black curve in Fig 1) and with the new model with high contact spheres and with toes (**B**; dashdot orange curve in Fig 1). We can appreciate the difference in knee flexion during early stance. The ground reaction force vector passes close to the knee joint center during early stance for the model without toes (**C**), whereas it is posterior to the knee joint center for the model with toes (**D**), requiring knee extension torques.

### Influence of the toe joints

Predictive simulations with a model incorporating toe joints produced stance knee flexion angles, knee extension torques, and ankle flexion torques that better reproduced experimental data as compared to simulations produced with a model with locked toe joints (Figs 1–2; results for all joints and muscles are shown in S1-3 Figs; animations are shown in S1-2 Movies). The predicted ankle kinematics changed when incorporating toe joints but did not qualitatively improve when compared to experimental data. Ground reaction forces produced with models with and without toe joints were comparable, although incorporating toe joints decreased the first peak of the vertical forces. In models with toe joints, the knee extensor (vasti) activations slightly increased during early stance, whereas the triceps surae (gastrocnemius and soleus) activations decreased during stance. The metabolic cost of transport was ~6 (7)% higher with the new (old) model with toe joints (4.18 J kg^−1^ m^−1^ for both old and new models) as compared to the new (old) model without toe joints (Table 1).

Interestingly the model adjustments detailed above (mass distribution and vertical location of the contact spheres) had barely any effect when incorporating toe joints in the model. In more detail, predictive simulations with the old model with a heavier torso and lighter legs, with low contact spheres, and without toe joints were almost identical as those produced with the new model with a lighter torso and heavier legs, with high contact spheres, and with toe joints (Fig 1).

### Influence of the Achilles tendon stiffness

Decreasing the Achilles tendon stiffness by more than 20% had a large effect on the knee and ankle kinematics during stance, but overall did not improve the realism of the simulations (Fig 3). The predicted knee torques were also affected by the change in tendon stiffness but the predicted ankle torques barely changed. Soleus activity increased with decreasing tendon stiffness. Furthermore, the soleus activation profile switched pattern when the decrease in tendon stiffness was larger than 20%, with the soleus turning on earlier with decreasing tendon stiffness. The gastrocnemius activity barely changed when adjusting tendon stiffness. For both muscles, the normalized fiber length profile was only slightly affected by the change in tendon stiffness, although both muscles operated at lower normalized fiber lengths with decreasing tendon stiffness. Metabolic cost of transport and stride length increased with decreasing tendon stiffness.

**Fig 3:**
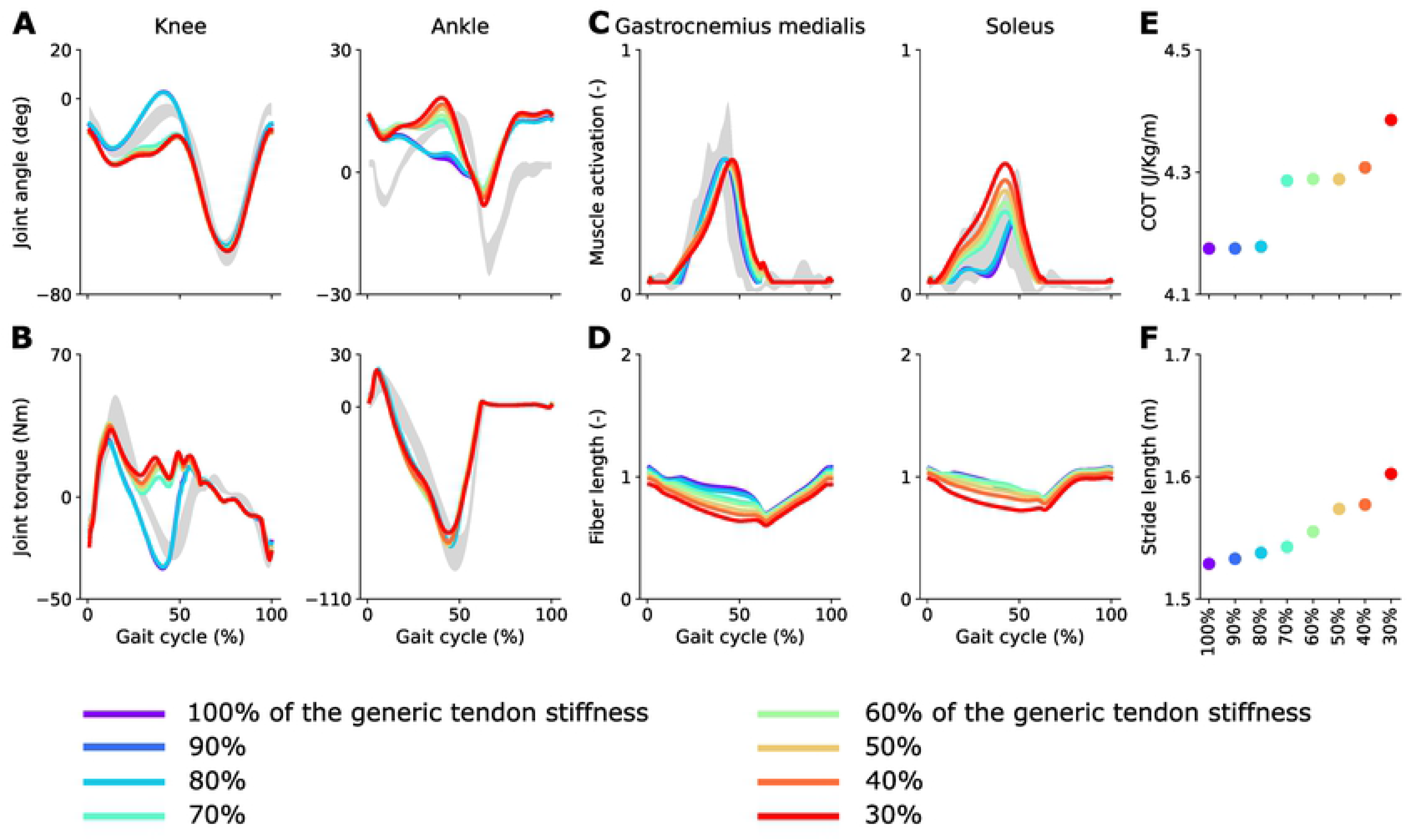
Influence of Achilles tendon stiffness on predicted gait pattern. Predicted knee and ankle kinematics (**A**) and kinetics (**B**), triceps surae activations (**C**), and normalized muscle fiber lengths (**D**) with models with different Achilles tendon stiffness. Predicted metabolic cost of transport (**E**) and stride length (**F**) as a function of the Achilles tendon stiffness (in percent of the nominal value). Experimental data (shaded areas) are shown as mean ± 2 standard deviations. The experimental electromyography data were normalized to peak nominal activations (100% of the generic Achilles tendon stiffness).

### Influence of the convergence tolerance and mesh density

Increasing the number of mesh intervals per half gait cycle had barely any influence on the results (i.e., state and control trajectories) obtained with the models with and without toe joints (Table 2), suggesting that simulation accuracy was good. When selecting the lowest optimal cost value from both initial guesses, the optimal cost value decreased (i.e., improved) by only about 1.4% and 1.3% between 50 and 125 mesh intervals for simulations produced with models with and without toe joints, respectively. When we used only 50 mesh intervals, both initial guesses led to different solutions. In particular, simulations from the cold-start had a much higher optimal cost value (464.7 vs 373.3 and 344.6 vs 290.7 for models with and without toe joints, respectively), meaning that the optimization algorithm converged to different local minima. Increasing the mesh density resulted in similar solutions for both initial guesses. This was observed for models with and without toe joints. Overall, the number of iterations increased when increasing the mesh density, but with no clear pattern. At high mesh density (100 and 125), the number of iterations was lower for the cold-start than for the hot-start.

**Table 2:** Influence of the mesh density on the convergence profile.

|  |  | N=50 |  | N=75 |  | N=100 |  | N=125 |  |
| --- | --- | --- | --- | --- | --- | --- | --- | --- | --- |
|  |  | # Iter | Cost | # Iter | Cost | # Iter | Cost | # Iter | Cost |
| With<br>toe<br>joints | Hot-start | 795 | 373.3 | 2541 | 371.2 | 1359 | 369.7 | 2052 | 368.1* |
|  | Cold-start | 1047 | **464.7** | 1363 | 369.4 | 1299 | 369.7 | 1232 | 368.3 |
| Without<br>toe<br>joints | Hot-start | 917 | 290.7 | 969 | 288.4 | 1860 | 287.3 | 2097 | 286.7 |
|  | Cold-start | 1012 | **344.6** | 1197 | 288.3 | 894 | 287.5 | 988 | 287.0* |

N is the number of mesh intervals, # Iter is the number of iterations before convergence, and Cost is the optimal cost value (integral of equation 1 with the optimal values). Bold values indicate large differences between both initial guesses and \* indicate solutions with the lowest optimal cost values.

Tightening the convergence tolerance had barely any influence on state and control trajectories produced with models with and without toe joints (Table 3), except for small differences in shoulder rotation. Specifically, when selecting the lowest optimal cost value from both initial guesses, the optimal cost value decreased by less than 0.1% when lowering the convergence tolerance from 10^−4^ to 10^−6^, suggesting that a convergence tolerance of 10^−4^ is strict enough. On average, the number of iterations was about twice and thrice as large for a convergence tolerance of 10^−5^ vs 10^−4^ and 10^−6^ vs 10^−4^, respectively.

**Table 3:** Influence of the convergence tolerance on the convergence profile.

| | | Tolerance = $1e^{-4}$ | | Tolerance = $1e^{-5}$ | | Tolerance = $1e^{-6}$ | |
| --- | --- | --- | --- | --- | --- | --- | --- |
|  |  | # Iter | Cost | # Iter | Cost | # Iter | Cost |
| With<br>toe<br>joints | Hot-start | 1359 | 369.7 | 2406 | 369.5 | 4452** | 369.5* |
|  | Cold-start | 1299 | 369.7 | 2559 | 369.5 | 4812 | 369.5 |
| Without<br>toe<br>joints | Hot-start | 1860 | 287.3 | 2894 | 287.2 | 3820 | 287.2* |
|  | Cold-start | 894 | 287.5 | 2782 | 287.2 | 3753 | 287.2 |

Tolerance refers to the IPOPT convergence tolerance, # Iter is the number of iterations before convergence, and Cost is the optimal cost value (integral of equation 1 with the optimal values). \* indicate solutions with the lowest optimal cost values. \*\* indicates a solution that was not optimal and for which the IPOPT restoration phase failed.

## Discussion

Musculoskeletal modeling choices can have a large influence on predictive simulations of walking. Here, we investigated the effect of two out of many possible modeling parameters, namely the number of degrees of freedom in the foot and the stiffness of the Achilles tendon, inspired by two common shortcomings of predictive simulations: a lack of stance knee flexion and terminal ankle plantarflexion. These gait features likely result from the complex interplay between motor control and musculoskeletal dynamics. While previous studies have focused on the effect of the performance criterion on the predicted walking pattern [4,10], our study reveals that a more physiologically plausible model of the feet, which incorporate toe joints, contributes to robustly eliciting knee flexion during stance. The Achilles tendon stiffness also had a considerable influence on the simulated gait pattern, but decreasing it did not improve the realism of the simulations. We argue that the causal relationships between musculoskeletal properties and gait patterns that this study revealed are not intuitive, underlining the power of computer simulations to shed lights on the role of isolated neuro-musculoskeletal features in shaping walking patterns. Yet, given that many combinations of performance criteria and musculoskeletal properties might generate the same gait features, experimental validation remains critical to ascertain that simulations are physiologically plausible. While it is clear that humans have toes, determining muscle-tendon properties and geometry experimentally is more challenging.

Models of foot-ground interaction shape how ground reaction forces are transmitted through the joints, which might explain why adding toe joints had a large influence on the more proximal knee joints. We also found that the position of the contact spheres can affect gait features such as stance knee flexion angles and knee extension torques, which further supports the importance of foot-ground interaction models. In comparison to our previous simulation study [10], we adjusted the vertical position of the contact spheres such that the model better captured the experimentally observed distance between the ankles and the ground. This change in the ground contact model, together with the altered mass distribution over the segments, explains why the simulations based on the model without toes presented here have larger stance knee flexion angles and knee extension torques than the simulations we previously published [10]. However, when incorporating toes the simulated gait patterns were robust against these changes in contact location and mass distribution (Fig 1). This agrees with our expectation that subjects wearing shoes with thicker soles and having a heavier trunk would still walk with stance knee flexion. The current foot and foot-ground contact models still overly simplify the complex human foot and foot-ground interactions. Simulations could help further elicit the role of foot dynamics in shaping walking patterns.

Although our results suggest that toes contribute to eliciting stance knee flexion, motor control objectives might also shape stance knee flexion. Previous simulation studies have demonstrated that the performance criterion influences stance knee flexion (e.g., [4]). Yet the relationship between the performance criterion and stance knee flexion has not been unambiguous, suggesting that other factors play a role as well. Other performance criteria, such as reducing impact and improving stability [7], have also been suggested to induce stance knee flexion but, as far as we know, these hypotheses have not been tested either in simulation or through experiments. While it would be technically straightforward to penalize impact in the cost function when performing predictive simulations, it is more challenging to incorporate stability constraints. Simulations can be considered stable when they are robust against internal and external perturbations. Stability requires feedback control to react to unexpected perturbations, which would require solving stochastic optimal feedback simulations. Whereas such robust simulations have been generated for simple models [31], it remains challenging to perform robust simulations based on complex models. Predictive simulations of walking relying on reflex-driven control models have been shown to be robust against perturbations [32]. Yet predicting realistic knee mechanics has also been challenging with such models [5,33,34], possibly because the pre-imposed reflex structure does not sufficiently capture human motor control or because of model simplifications such as the absence of toe joints. In this study, we used the same cost function for all simulations, and tested the influence of musculoskeletal modeling assumptions. Studying the interactions between mechanics and cost function was out of the scope of this work, but should be envisioned in future studies. Indeed, it is now difficult to draw general conclusions, since the effect of the mechanics might be different when using a different performance criterion.

We found that incorporating toe joints in the model increased the simulated cost of transport, suggesting that having toes does not improve locomotion efficiency. The increased metabolic energy expenditure in the knee extensors is in line with previous observations that knee flexion during stance is energetically costly [4]. Yet our simulations suggest that, with toes, a gait pattern requiring knee extension torques during stance optimizes performance, possibly because, as we found, it reduces the metabolic energy expenditure in the plantarflexors with respect to a straight knee pattern in the absence of toes. Song et al. also found that foot compliance increased metabolic cost taking a different approach. Instead of only adding toes, they also modeled compliance of the foot arch and plantar fascia and used reflex-driven 2D simulations of walking [35]. Our simulation results are in line with experimental observations during running where using shoe wear that increases foot stiffness reduces metabolic cost [36].

Our simulations with different Achilles tendon stiffness suggest that performance optimization might encourage muscles to work at relatively low contraction velocities. Such muscle behavior agrees with experimental studies showing that the fascicles of the gastrocnemius tend to act relatively isometrically during the stance phase of walking [37]. While decreasing Achilles tendon stiffness by more than 20% had a big influence on ankle and knee kinematics, plantarflexor muscle fiber lengths changed more gradually with tendon stiffness and ankle plantarflexor torques were insensitive to tendon stiffness. Decreasing Achilles tendon stiffness improved terminal stance ankle plantarflexion but this came at the cost of less realistic knee angles and torques. A possible explanation for our observations is that the relationship between joint kinematics and plantarflexor muscle-tendon lengths is inaccurately modeled. Indeed, fiber length does not only depend on tendon stiffness but also on muscle-tendon length. If we overestimate the decrease in muscle-tendon lengths with ankle plantarflexion, this would result in reduced ankle plantarflexion in predictive simulations where having nearly isometric fiber lengths seems to optimize performance. This hypothesis is supported by previous results from inverse analysis comparing simulated fiber lengths to their experimental counterparts measured using ultrasound [38]. In that study, we found that when imposing joint kinematics and kinetics, the simulated change in gastrocnemius fiber length in terminal stance exceeded the measured change. In addition, previous studies have reported a wide range of Achilles tendon moment arms, and moment arms determine how muscle-tendon lengths change with joint angles [39–41]. We therefore believe that more accurately modeling plantarflexor geometry might improve simulated ankle angles.

Our analysis of convergence suggests that the selected discretization (i.e., number of mesh intervals) is accurate enough, the selected convergence tolerance is strict enough, and the solutions are robust against different initial guesses. Overall, this suggests that the control and state trajectories were approximated with sufficient accuracy. Note that this does not mean that our results are not local optima. It is indeed possible that both straight and flexed knee patterns during stance are local optima and that musculoskeletal modeling assumptions make one local optima more likely to be found that the other one. Our results suggest the importance of comparing simulations from different initial guesses, since they may be completely different (e.g., hot-start vs cold-start in simulations with 50 mesh intervals; Table 2). Ensuring that simulations are not impacted by the use of a finer mesh, a tighter convergence tolerance, or the choice of the initial guess brings is important to gain confidence in the accuracy of the results. It was interesting, although surprising, to note that our simulations converged faster (i.e., with fewer iterations) from the cold-start than from the hot-start when using a high mesh density. This suggests a limited impact of using a hot-start for such simulations, but this observation should be confirmed in future studies before generalizing. Finally, while our results suggest that we could have run our simulations using a convergence tolerance of 10^−4^ and 50 mesh intervals per half gait cycle, we cannot ensure that such numerical choices will be valid when altering musculoskeletal mechanics or cost function, or when simulating different movements.

## Conclusion

Both mechanics and cost function shape simulations of human walking. While previous studies have focused on the role of the cost function, we here demonstrate the effect of mechanical assumptions on predictive simulations of walking. Incorporating toes in the musculoskeletal model contributed to robustly eliciting stance knee flexion. Yet it did not improve terminal stance ankle plantarflexion, nor did decreasing the stiffness of the Achilles tendon. The lack of ankle plantarflexion at terminal stance in predictive simulations of walking thereby remains an open question, and further work is required to better understand how mechanical factors shape human walking. Computer simulations provide a useful tool to investigate the effect of the mechanics on human movements. However, as model complexity increases, so do the interactions between modeling assumptions, making it more difficult to comprehensively assess the effect of model parameters. Validating through experiments the effect of modeling assumptions observed in simulations would increase confidence in simulation outcomes, contributing to their deployment for applications such as personalized medicine and assistive device design.

## Acknowledgments

The authors would like to thank Tom Van Wouwe for helping with data collection.

## Supporting information

**S1 Fig. Predicted joint kinematics.**

Experimental data (shaded areas) are shown as mean ± 2 standard deviations. The standard deviation for the pelvis tz (lateral pelvis displacement) is not shown due to large variations caused by different walking directions followed by the subject during the data collection. The old and new models have the same mass but different mass distribution: the new model having a lighter torso but heavier legs as compared to the old model.

**S2 Fig. Predicted joint kinetics.**

Experimental data (shaded areas) are shown as mean ± 2 standard deviations.

**S3 Fig. Predicted muscle activations.**

Experimental data (shaded areas) are shown as mean ± 2 standard deviations. The experimental electromyography data were normalized to peak activations of the new model with high contact spheres and with toes (dashdot orange curve).

**S1 Movie. Predictive simulations of walking with the old model (heavier torso but lighter legs) with low contact spheres and without toe joints.** Muscles turn red when active. The green arrows represent the ground reaction forces. The playback speed is 0.2 times real-time.

**S2 Movie. Predictive simulations of walking with the new model (lighter torso but heavier legs) with high contact spheres and with toe joints.** Muscles turn red when active. The green arrows represent the ground reaction forces. The playback speed is 0.2 times real-time.

